# Natural gene variation in *Cannabis sativa* unveils a key region of cannabinoid synthase enzymes

**DOI:** 10.1101/2023.08.30.555511

**Authors:** Cloé Villard, Christian Bayer, Nora Pasquali Medici, Arjen C. van de Peppel, Katarina Cankar, Francel Verstappen, Iris F. Kappers, M. Eric Schranz, Bastian Daniel, Robin van Velzen

## Abstract

Cannabinoids are well-known specialised metabolites from the plant *Cannabis sativa* L. (cannabis). They exhibit various therapeutical to intoxicating psychoactive effects and have potential for medicinal applications. Among the enzymes involved in cannabinoid biosynthesis, cannabinoid oxidocyclases such as the tetrahydrocannabinolic acid (THCA) synthase play a key role in determining cannabis chemotype. To improve our understanding of cannabinoid oxidocyclase structure-function relationship, we proposed a new approach to targeted mutagenesis. By reviewing cannabis natural variation, three cannabinoid oxidocyclase mutations (S355N, CONF, G376R) associated to atypical plant chemotypes were selected. *In-vitro* characterization of THCA synthase mutants demonstrated these mutations significantly impact enzyme activity, correlating with the associated chemotype: S355N nearly inactivated the THCA synthase, CONF impaired CBGA metabolization and altered product specificity, while G376R drastically reduced enzyme activity and altered product specificity. *In-silico* docking experiments permitted to model the successive steps of THCA synthase substrate metabolization, revealing that the three mutations hamper substrate binding. Collectively, our results demonstrated how plant diversity can be leveraged to guide enzyme targeted mutagenesis, highlighted a key region of cannabinoid oxidocyclases, and permitted the establishment of a new model of the THCA synthase catalytic mechanism. This provides new insights into enzyme function, which can ultimately help developing medicinal cannabis cultivars and cannabinoid biotechnological production.

## Introduction

*Cannabis sativa L.* (cannabis) is a well-known plant from the Cannabaceae family which exhibits therapeutic and intoxicating psychoactive properties. These properties come from terpenophenolic specialized metabolites called cannabinoids (Gaoni and Mechoulam, 1971; Mechoulam, 2005; Casajuana Köguel et al., 2018). Cannabinoids are synthesized in cannabis glandular trichomes as carboxylic acids that do not have psychoactive effects (Sirikantaramas et al., 2005; Livingston et al., 2020). However, upon exposure to light or high temperature (*i.e.*, smoking, baking), cannabinoid acids are non-enzymatically decarboxylated into neutral cannabinoids (**Figure 1A**) exhibiting psychoactive activities (Sirikantaramas et al., 2004; Wang et al., 2016; ElSohly et al., 2017). So far, more than 120 cannabinoids have been identified in cannabis, the most abundant and best-known being Δ9-tetrahydrocannabinolic acid (THCA), cannabidiolic acid (CBDA), and their neutral counterparts Δ9-tetrahydrocannabinol (THC) and cannabidiol (CBD) (ElSohly et al., 2017). At high doses, THC is intoxicating (Lupica et al., 2004), but at low dosage, it has therapeutic potential such as analgesic, appetite or sleep stimulant (Costa, 2007; van de Donk et al., 2019; Suraev et al., 2020). CBD is not intoxicating, modulates THC activity, and exhibits medicinal properties such as anti-inflammatory, anti-nausea, anti-anxiety or anti-arthritic (Burstein, 2015; Boggs et al., 2018; Casajuana Köguel et al., 2018; Colizzi et al., 2020).

**Figure 1.**
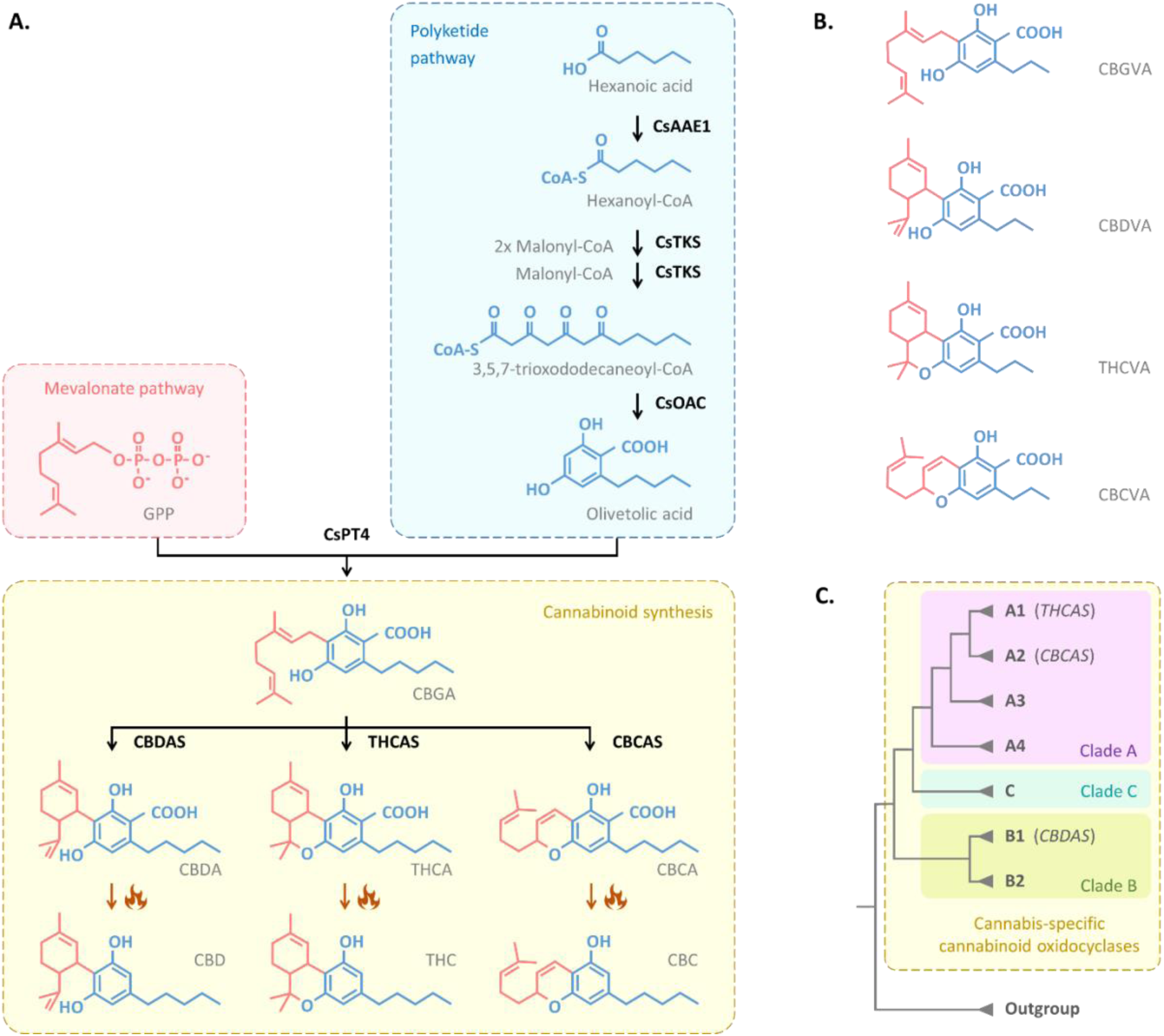
Production of cannabinoids in *Cannabis sativa*. **(A)** Biosynthesis pathway of pentyl-cannabinoids. Enzymatic reactions are symbolized with black arrows, nonenzymatic decarboxylations with dark orange arrows and heat symbols (Taura et al., 1995; Taura et al., 1996; Fellermeier and Zenk, 1998; Morimoto et al., 1998; Sirikantaramas et al., 2004; Taura et al., 2007; Taura et al., 2009; Stout et al., 2012; Gagne et al., 2012; Laverty et al., 2019; Luo et al., 2019). **(B)** Propyl-cannabinoid acid structures. **(C)** Phylogenetic-based classification of cannabinoid oxidocyclases, based on van Velzen and Schranz (2021). Main clades are identified with colored blocks; subclades are written in grey. Abbreviated molecules: CBC, cannabichromene; CBCA, cannabichromenic acid; CBCVA, cannabichromevarinic acid; CBD, cannabidiol, CBDA, cannabidiolic acid; CBDVA, cannabidivarinic acid; CBGA, cannabigerolic acid; CBGVA, cannabigerovarinic acid; GPP, geranyl pyrophosphate; THC, Δ9-tetrahydrocannabinol; THCA, Δ9-tetrahydrocannabinolic acid; THCVA, Δ9-tetrahydrocannabivarinic acid. Abbreviated enzymes: CsAAE1, *C. sativa* acyl activating enzyme 1; CsTKS, *C. sativa* tetraketide synthase; CsOAC, *C. sativa* olivetolic acid cyclase; CsPT4, *C. sativa* prenyltransferase 4; CBDAS, CBDA synthase; THCAS, THCA synthase; CBCAS, CBCA synthase.

The biosynthesis pathway of THCA and CBDA in cannabis has been fully elucidated: the genes and enzymes catalysing the successive reactions were identified and characterized (**Figure 1A**), making it possible to produce cannabinoids from simple sugars in engineered microorganisms (Luo et al., 2019). Briefly, cannabinoid biosynthesis starts in the polyketide pathway with the production of olivetolic acid (Taura et al., 2009; Stout et al., 2012; Gagne et al., 2012) and its prenylation into the central precursor cannabigerolic acid (CBGA) (Fellermeier and Zenk, 1998; Luo et al., 2019). CBGA can then be converted into CBDA, THCA or other derived cannabinoid acids such as cannabichromenic acid (CBCA), through stereospecific oxidative cyclizations catalysed by cannabinoid oxidocyclases (Taura et al., 1995; Taura et al., 1996; Morimoto et al., 1998; Sirikantaramas et al., 2004; Taura et al., 2007; Laverty et al., 2019). In addition to the above-mentioned cannabinoids which possess a pentyl side chain, cannabis also produces cannabinoids from divarinolic acid, an olivetolic acid variant with a propyl side chain. This leads to the propyl-variants cannabigerovarinic acid (CBGVA), Δ9-tetrahydrocannabivarinic acid (THCVA), cannabidivarinic acid (CBDVA) and cannabichromevarinic acid (CBCVA, **Figure 1B**). The biosynthesis of propyl-cannabinoids has been less studied, but is assumed to be mediated by the same enzymes producing pentyl-cannabinoids (De Meijer and Hammond, 2016; Luo et al., 2019).

The last step of cannabinoid biosynthesis, branching point to molecules with different bioactivities, is of particular interest. So far, three cannabinoid oxidocyclases have been characterized: the CBDA, THCA and CBCA synthases (CBDAS, THCAS, CBCAS) which convert CBGA into either CBDA, THCA and CBCA, respectively, as major products (Sirikantaramas et al., 2004; Taura et al., 2007; Laverty et al., 2019). As minor products, the THCAS and CBDAS enzymes also yield some CBCA, CBDA or THCA, and other unidentified cannabinoids (Zirpel et al., 2018). The *THCAS* and *CBCAS* genes are 1638-bp intronless open reading frames (ORF) sharing 96% sequence identity (Sirikantaramas et al., 2004; Laverty et al., 2019). The *CBDAS* gene comprises a 1635-bp intronless ORF sharing 89% identity with *THCAS* and *CBCAS* (Taura et al., 2007). Many *CBDAS*, *THCAS*, *CBCAS* variants and other uncharacterized cannabinoid oxidocyclase genes have been reported, *e.g.*, Onofri *et al*. (2015), Cascini *et al*. (2019), Fulvio *et al*. (2021). To categorize these sequences, van Velzen and Schranz (2021) proposed a phylogenetic-based classification which describes 3 main clades (A, B, C) subdivided into 7 subclades (A1-4, B1-2, C) (**Figure 1C**). The *THCAS*, *CBCAS* and *CBDAS* correspond to subclades A1, A2 and B1, respectively, while subclades A3, A4, B2 and C contain pseudogenes and uncharacterized genes.

Cannabinoid oxidocyclases are members of the larger berberine bridge enzyme (BBE)-like (Sirikantaramas et al., 2004; Taura et al., 2007), a family of oxidoreductases known to catalyze a variety of chemically challenging reactions (Daniel et al., 2017). Like other BBE-like, cannabinoid oxidocyclases are bi-covalently bound to a flavin adenine dinucleotide (FAD) cofactor and oxidize their substrate by reducing FAD (Sirikantaramas et al., 2004; Winkler et al., 2009; Shoyama et al., 2012). Targeted mutagenesis studies (Sirikantaramas et al., 2004; Taura et al., 2007; Zirpel et al., 2018) and the elucidation of THCAS crystal structure (Shoyama et al., 2012) offered valuable insights into cannabinoid oxidocyclase activity, allowing the identification of some residues involved in FAD binding, substrate binding, catalysis, disulfide bridge, and glycosylation (**Supplemental Table S1**), and to propose a catalytic mechanism for the conversion of CBGA into THCA (Shoyama et al., 2012). However, all mutagenesis studies conducted so far focused on artificially designed mutants. As a result, we do not know yet how mutations in these enzymes affect cannabis chemotype. The only exception is our previous study, van Velzen *et al*. (unpub.), in which we identified a natural THCAS substitution in cannabis plants with [atypical chemotype, CONF] and demonstrated that this mutation lower THCA production.

In this study, we propose a new targeted mutagenesis approach, based on plant natural variation, to improve our understanding of cannabinoid oxidocyclase structure-function relationship. Through reviewing cannabinoid oxidocyclase sequence variation and associated plant chemotypes, we identified candidate amino acid substitutions. *In-vitro* site-directed mutagenesis and *in-silico* docking experiments revealed these substitutions have a significant negative impact on substrate binding and therefore THCAS activity and allowed us to propose a new model for THCAS catalytic mechanism. This led us to identify a new critical region within cannabinoid oxidocyclase enzymes, which can affect plant chemotype.

## Results

### Using natural variation to identify substitutions of interest

Reviewing previously published genes, we compiled a dataset containing about 200 cannabinoid oxidocyclase sequences and categorized them according to the most recent classification (van Velzen and Schranz, 2021). Sequences from the THCAS (A1), CBCAS (A2) and CBDAS (B1) subclades were individually screened, and a total of 36, 54 and 17 single nucleotide polymorphisms (SNP) involving amino acid (aa) substitutions were identified, respectively (**Supplemental Table S2**). Seven substitutions were similar in *THCAS* and *CBCAS* or in *CBCAS* and *CBDAS* sequences, leaving a total of 100 different aa substitution positions (**Supplemental Table S2**). To provide structural information and a context around these substitution positions, they were visualized on the THCAS three-dimensional (3D) model and compared with previously studied residues (**Figure 2A, 2B, Supplemental Table S1**; Sirikantaramas et al., 2004; Taura et al., 2007; Shoyama et al., 2012; Zirpel et al., 2018; van Velzen et al., unpub.). Note that, compared to THCAS and CBCAS, the CBDAS enzyme possesses a gap at codon position 253. We have therefore numbered CBDAS residues (n ≥ 253) according to their homologous position in THCAS/CBCAS (n+1; altered numbers are highlighted with an asterisk). 22 substitution positions were relatively close to the active site. The others were on the surface of the enzyme and/or far from known regions of interest (**Figure 2, Supplemental Table S2**).

**Figure 2.**
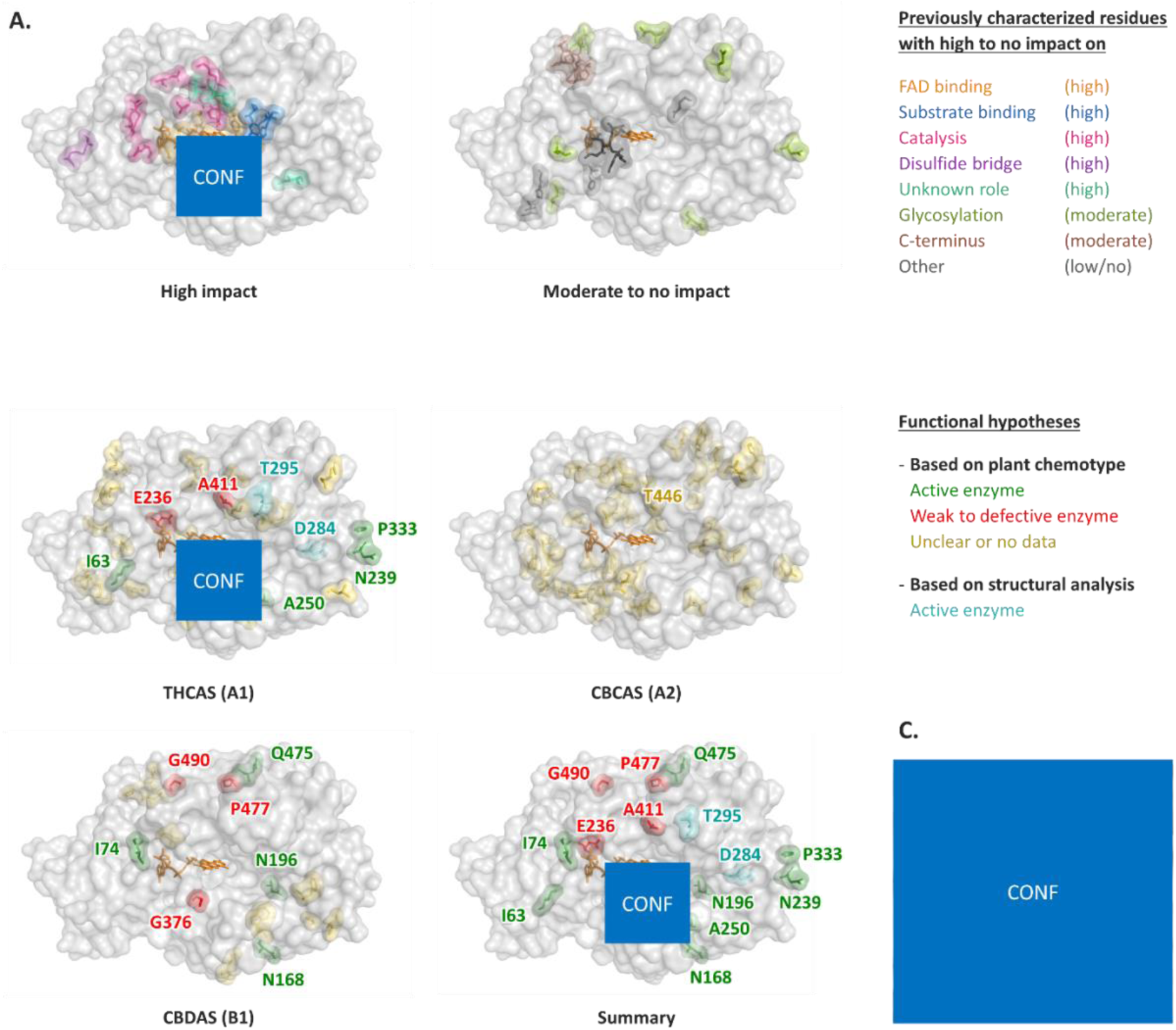
Cannabinoid oxidocyclase residues of interest, visualized in the THCAS 3D model. The FAD cofactor is represented in orange. **(A)** Residues inquired in previous cannabinoid oxidocyclase site-directed mutagenesis studies or described in the elucidation of the THCAS crystal structure, colored according to their functional role and impact on enzyme activity (**Supplemental Table S1**). **(B)** Residues for which aa substitutions have been identified in THCAS, CBCAS and CBDAS sequences. Residues are colored according to their hypothesized impact on enzyme activity, based on chemotype data (Kojoma et al., 2006; McKernan et al., 2015; Onofri et al., 2015; Weiblen et al., 2015; Zirpel et al., 2018; Cascini et al., 2019; Gao et al., 2020; Garfinkel et al., 2021; Fulvio et al., 2021; van Velzen et al., unpub.) or structural analysis (Ali et al., 2019; **Supplemental Table S2**). **(C)** Zoom on the residues S355, CONF and G376.

To provide functional hypotheses about the aa substitutions, we then relied on plant chemotype, when available in the literature (**Table 01**, **Supplemental Table S2**). On the one hand, four THCAS (I63L, A250D, N329T, P333R) and four CBDAS substitutions (T74S, N168S, N196S, K475Q*) were identified in plants producing normal levels of THCA and CBDA, respectively (Kojoma et al., 2006; McKernan et al., 2015; Onofri et al., 2015). Since these plants necessarily possess active THCAS and CBDAS, authors hypothesized the substitutions to have low to no impact on enzyme activity (**Table 01**). Interestingly, three THCAS substitutions (D284V, T295P, K377Q) identified in plants of unknown chemotype (Ali et al., 2019; Gao et al., 2020) were also associated to low functional impact by Ali *et al*. (2019) who performed structural bioinformatic analyses (D284V, T295P) and Zirpel *et al*. (2018) through site-directed mutagenesis (K377Q, **Supplemental Table S1**). Nine of the eleven substitutions associated to low functional impact affected residues on the surface of the enzyme or relatively far from the active site (**Table 01**, **Figure 2B**).

On the other hand, three THCAS (E236Q, S355N, A411V) and three CBDAS substitutions (G376R*, P477S*, G490R*) originated from plants accumulating CBGA and/or with unusually low THCA and CBDA levels (McKernan et al., 2015; Onofri et al., 2015; Garfinkel et al., 2021). Authors therefore hypothesized that these substitutions weaken or inactivate cannabinoid oxidocyclases (**Table 01**). Two other residues (CONF, T446) were associated to high functional impact through site-directed mutagenesis experiments: the CONF substitution was found in plants with [atypical chemotype, CONF] and demonstrated to reduce the THCAS activity (van Velzen et al., unpub.). The T446A CBCAS substitution was identified in cannabis of unknown chemotype (Weiblen et al., 2015), but Zirpel *et al*. (2018) demonstrated that the artificial T446I THCAS and I446T* CBDAS mutations weakened the enzyme activity (**Supplemental Table S1**). Six of these eight substitutions associated with high functional impact were located close to the active site (**Figure 2B**, **Table 01**). Interestingly, the S355N and G376R* substitutions were close to CONF (**Figure 2C**), and all three residues are located in a looped region adjacent to the active site extending from Y354 to A380 (ASA-loop). This intriguing pattern motivated us to investigate this region by (further) characterizing the impact of substitutions S355N (plants accumulating CBGA), CONF (plants with atypical chemotype) and G376R* (plants with low CBDA level).

Finally, three THCAS (V125L, E265G, G410E) and eight CBDAS substitutions (H143R, F258S*, G319V*, V321M*, W347R*, V521A*, L540Q*, R542G*) were identified in plants of known cannabinoid content, but associated to unclear data (*i.e.*, multiple aa substitutions in the sequence or multiple alleles of the gene in the plant). For these and all remaining THCAS, CBCAS and CBDAS substitutions which had no associated chemotype data, no functional hypothesis could be formulated.

### Characterization of the S355N, CONF and G376R mutants

Residues S355, CONF and G376(*) are identical in the THCAS, CBCAS and CBDAS enzymes, and therefore probably play the same role in all three cannabinoid oxidocyclases. To generate comparable results, we tested the impact of mutations S355N, CONF and G376R in the THCAS. For this purpose, a wild-type (WT) and three single-mutant THCAS gene constructs were synthesized. Associated enzymes were expressed in yeast and used to perform *in-vitro* assays. In preliminary analyses, maximal WT THCAS activity was obtained at pH 5 and 50°C (data not shown), which is consistent with previous studies (Taura et al., 1995; Zirpel et al., 2018). While acid pHs are physiologically possible in plants, a temperature of 50°C is not the norm in cannabis, which is usually cultivated at about 25°C. Therefore, to test if the mutants were active, long incubations were performed at 50°C, but to ensure that the mutations could physiologically affect plant chemotypes, standardized assays were performed at 25°C.

Long incubations performed at 50°C showed that the CONF and G376R mutants, like the WT THCAS, could convert CBGA into THCA and CBCA (**Figure 3A**), and CBGVA into THCVA and CBCVA (**Figure 3B**). Trace amounts of CBDA and of an unknown compound assumed to be CBDVA were also detected. On the contrary, the S355N mutant did not convert CBGA into any detectable product, and incubation with CBGVA only yielded trace amounts of THCVA and CBCVA (**Figure 3A, 3B**).

**Figure 3.**
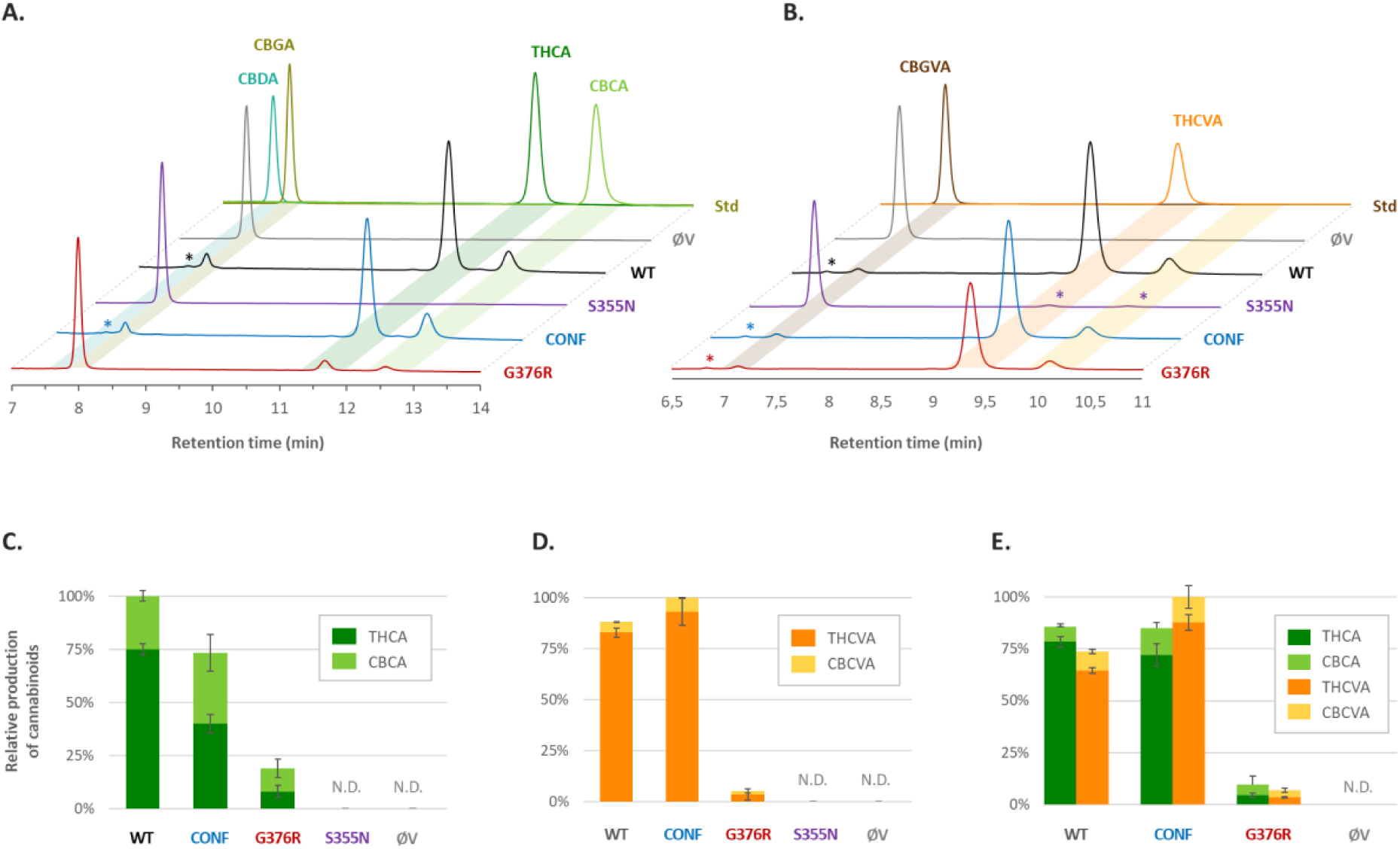
Metabolization of CBGA and its pentyl analog CBGVA by the WT and mutant THCAS enzymes. **(A, B)** Ultra-high-performance liquid chromatography (UHPLC) separation profiles. Enzymes were incubated at optimal 50°C in the presence of either CBGA **(A)** or CBGVA **(B)**. Cannabinoids were identified by comparison to standard molecules (lane Std). Peaks close to detection limit are highlighted with an asterisk (*). **(C, D, E)** Relative production of THC(V)A and CBC(V)A by the WT or mutant THCAS, incubated at physiological 25°C in the presence of CBGA **(C)**, CBGVA **(D)**, or equimolar CBGA and CBGVA **(E)**. An empty vector (ØV) was used as negative control. Incubations were performed in triplicates; error bars represent the standard deviation. N.D., not detected. Abbreviated molecules: CBCA, cannabichromenic acid; CBCVA, cannabichromevarinic acid; CBDA, cannabidiolic acid; CBGA, cannabigerolic acid; CBGVA, cannabigerovarinic acid; THCA, tetrahydrocannabinolic acid; THCVA, Δ9-tetrahydrocannabivarinic acid.

Standardized assays were then performed at 25°C with CBGA and/or CBGVA. When incubated with CBGA (**Figure 3C**), the CONF and G376R mutants displayed decreased activity, metabolizing respectively 72 ± 8 % and 19 ± 7 % of the CBGA metabolized by the WT. The CONF and G376R mutations also impacted THCAS product specificity toward a higher production of CBCA: the THCA/CBCA ratio of the WT enzyme was 3.0 ± 0.4 but was reduced to 1.3 ± 0.4 for the CONF mutant and 0.8 ± 0.1 for the G376R mutant (**Figure 3C**).

When incubated with CBGVA alone (**Figure 3D**), the activity of the CONF mutant was comparable to that of the WT (110 ± 13 %). The G376R mutant exhibited decreased activity, metabolizing only 6 ± 2 % of the CBGVA metabolized by the WT. The THCVA/CBCVA of the WT enzyme was 16.1 ± 0.9 but diminished to 13.4 ± 0.7 and 1.9 ± 0.5 for the CONF and G376R mutants, respectively (**Figure 3D**). For the S355N mutant, no product was detected. Although active and able to metabolize some CBGVA, the S355N mutant activity is therefore negligible.

Finally, the enzymes were incubated with equimolar concentrations of CBGA and CBGVA (**Figure 3E**). In these conditions, the WT showed a slight preference for CBGA, with a ratio CBGA_metabolized_/CBGVA_metabolized_ of 1.2 ± 0.1. This trend was reversed for the CONF mutant, which had a ratio of 0.9 ± 0.1. Substrate preference of the G376R mutant was less clear. Interestingly, incubations with both CBGA and CBGVA also affected the relative quantities of THC(V)A and CBC(V)A produced by the enzymes. In comparison to the incubations performed with CBGA alone, the THCA/CBCA ratio was about 3 to 4-fold higher for the WT (10.9 ± 2.1) and CONF mutant (5.8 ± 1.7). On the contrary, compared to the incubation performed with CBGVA alone, the THCVA/CBCVA ratio showed a 2-fold decrease for the WT (7.1 ± 0.8), CONF (7.2 ± 0.3) and G376R (1.1 ± 0.1) mutants (**Figure 3E**).

### Docking experiments helped decipher THCAS catalytic mechanism

To improve our understanding of THCAS activity, we next performed bioinformatic analyses. Despite the elucidation of THCAS crystal structure and the proposition of a catalytic mechanism for the conversion of CBGA into THCA (Shoyama et al., 2012), no 3D model including the substrate or product docked into the active site was available. We therefore performed *in-silico* docking experiments in the WT THCAS, aimed at identifying productive binding modes for the successive catalytic events corresponding to the conversion of CBGA into THCA. These events are, sequentially, (1) the activation of CBGA ionizable group by deprotonation, (2) the hydride transfer which forms an intermediate that can undergo cyclization, and (3), the intermediate pre-organization to form THCAS.

The main docking results are depicted in **Figure 4**. First, the CBGA substrate was docked into the WT THCAS enzyme (**Figure 4A**). The results showed that the CBGA carbon C2’ (*i.e.*, the atom undergoing oxidation) was located at 3.8 Å from the FAD position N5 (*i.e.,* the atom to which the hydride is transferred to). This short distance, together with the orientation of the hydrogen atoms and the respective angles of the CBGA-FAD complex are in perfect agreement with Fraaije and Mattevi (2000)’s concept of the site of oxidative attack, indicating a productive binding mode. In this configuration, the CBGA carboxy group was oriented toward H292 and Y417, two residues postulated to contribute to substrate binding via this polar interaction (**Supplemental Table S1**) (Shoyama et al., 2012). The CBGA prenyl-tail was located in a hydrophobic pocket of the THCAS, and was partly preorganized, *i.e.*, adopting a conformation similar to that of the final THCA product (compare **Figure 4A, 4D**). The CBGA C2 hydroxy group was oriented towards Y484, the THCAS most probable catalytic base (Shoyama et al., 2012) (**Supplemental Table S1**). Our dockings suggest that CBGA metabolization starts with the deprotonation of CBGA C2 hydroxy group, mediated by Y484. This deprotonation is followed by a hybrid transfer from CBGA C1’ to the FAD N5 position, leading to the formation of Intermediate I (**Figure 4B**). As the cyclization into THCA involves CBGA C4 hydroxy group, we postulate the existence of a second intermediate (**Figure 4C**). Intermediate II is formed via tautomerization of Intermediate I, involving the reprotonation of the C2 hydroxy group and simultaneous deprotonation of the conjugated C4 hydroxy group, with H_2_O (solvent) acting as a proton acceptor. The docking modes obtained for both Intermediates I (**Figure 4B**) and II (**Figure 4C**) are similar. Intermediate II is then preorganized. As it has the freedom to adopt THCA conformation, Intermediate II is finally cyclized into THCA, via a hetero-Diels-Alder reaction that forms a C6’C7’ double bond on the C4 oxygen (**Figure 4D**). We note that the conformation of both the aromatic moiety and the aliphatic olivetolic acid tail are consistent across the dockings of CBGA, both intermediates, and THCA.

**Figure 4.**
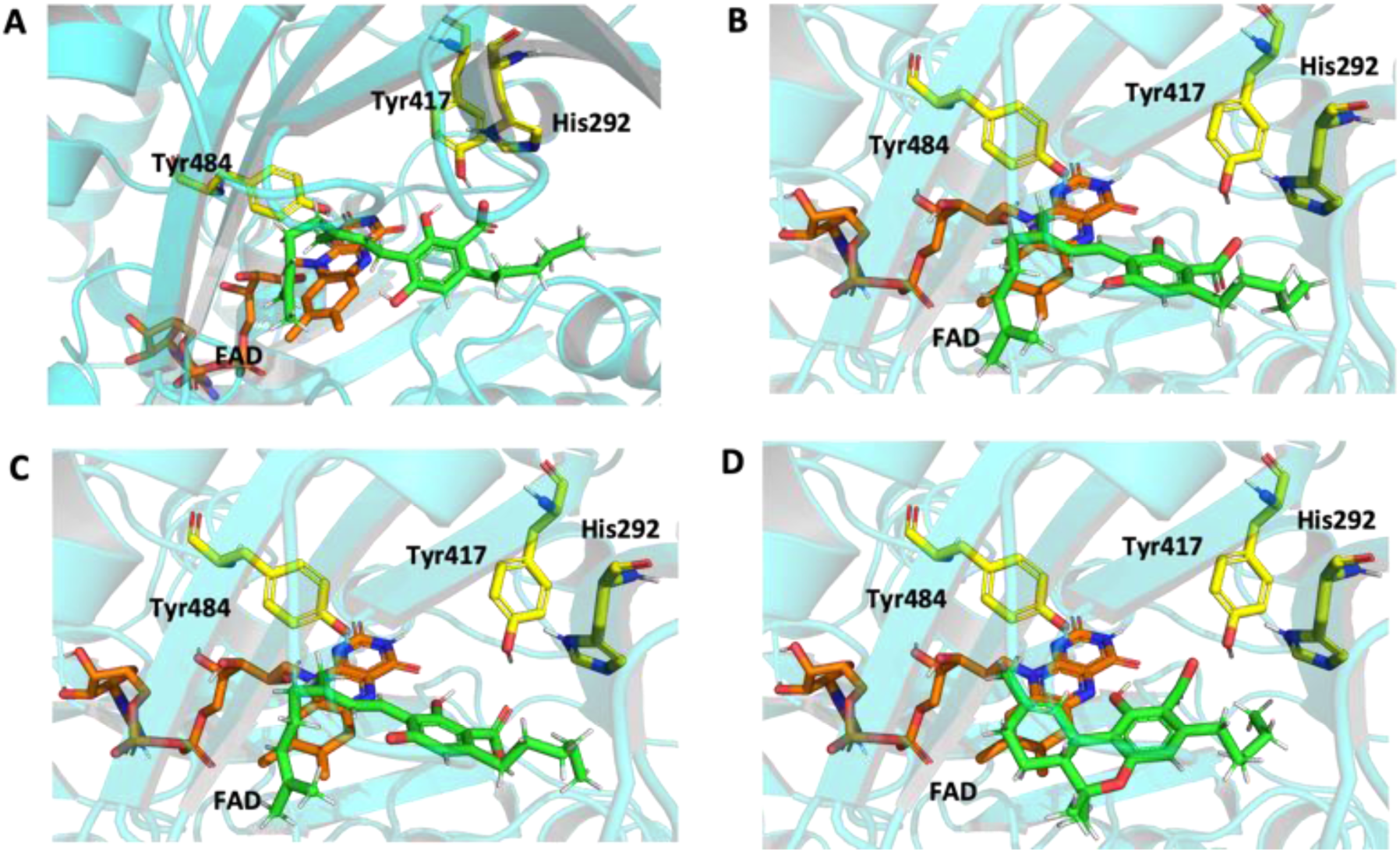
Molecular dockings of CBGA (A), intermediate I (B), intermediate II (C) and THCA (D) in the WT THCAS. The respective ligands are depicted in green, the FAD cofactor in orange, and catalytic relevant polar residues as yellow sticks. Abbreviated molecules: CBGA, cannabigerolic acid; FAD, flavin adenine dinucleotide; THCA, tetrahydrocannabinolic acid. Abbreviated enzyme: WT THCAS, wild type THCA synthase.

Based on these docking results, we propose the new updated mechanism corresponding to the conversion of CBGA into THCA, depicted in **Figure 5**. The WT THCAS can also convert CBGA into CBCA as minor product. We postulate that CBCA is formed via a similar hetero-Diels-Alder cyclization of Intermediate II, involving the C3′C2′ instead of the C7′C6′ double bond (mechanism not shown).

**Figure 5:**
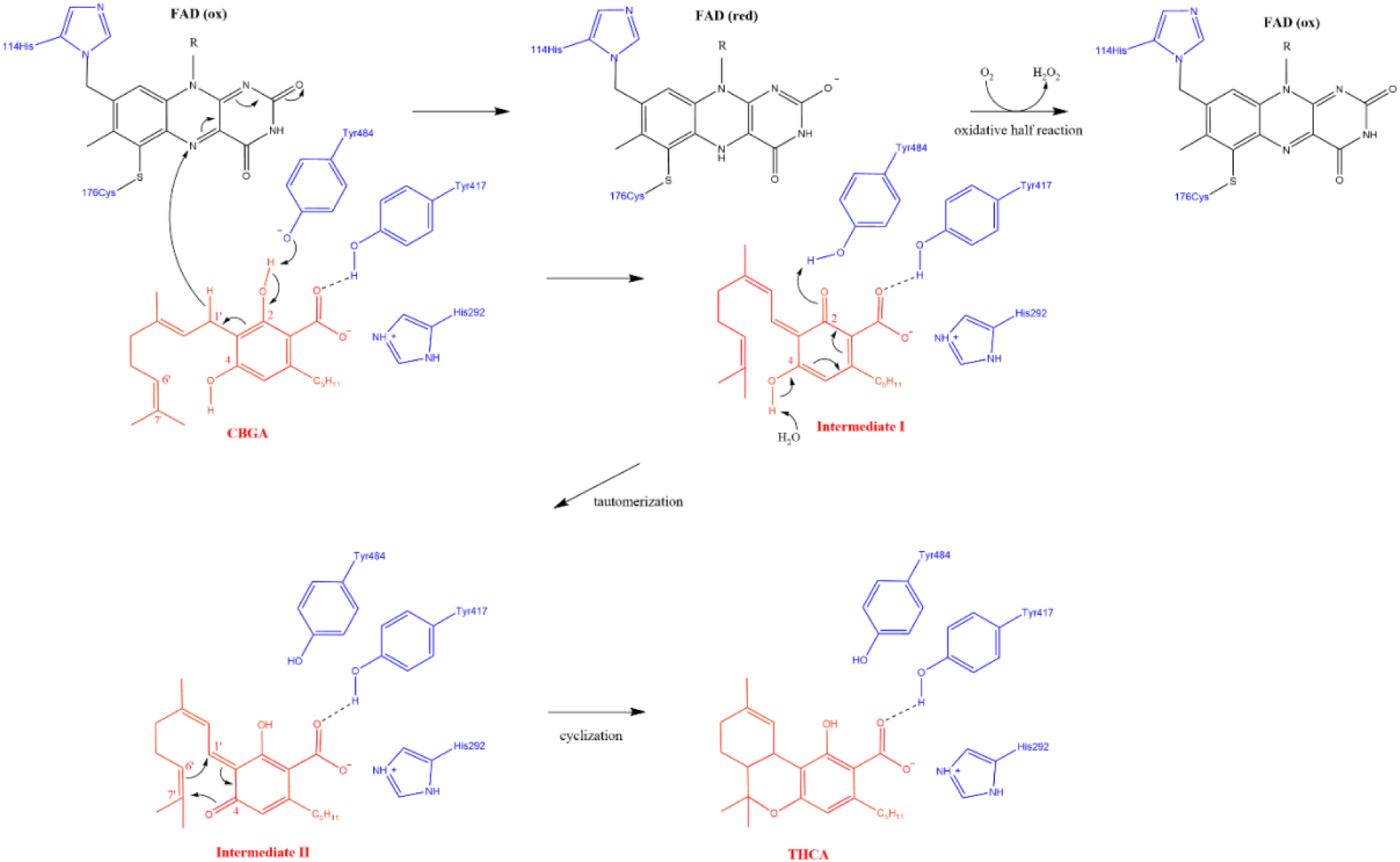
Updated WT THCAS catalytic mechanism. Catalytic residues are depicted in blue, the FAD co-factor in black, CBGA, the respective intermediates and THCA in red. Starting of the mechanism point is the deprotonation of the C2 hydroxyl group of CBGA by Y484 which serves as a catalytic base. Subsequently, a hydride transfer from the C1’ position to the N5 of FAD occurs, leading to the first Intermediate I. The oxygen next to the C2 position is then reprotonated and simultaneously the hydroxyl group next to the C4 position is deprotonated, where H_2_O functions as a proton acceptor. The resulting dienone Intermediate II then undergoes a hetero-Diels-Alder reaction. New bonds are formed between C6’ and C1’ and the C7’ and the C4 oxygen. Red, reduced; Ox, oxidated. Abbreviated molecules: CBGA, cannabigerolic acid; FAD, flavin adenine dinucleotide; THCA, tetrahydrocannabinolic acid. Abbreviated enzymes: WT THCAS, wild type THCA synthase.

### Bioinformatic analysis of the ASA-loop

To explain the impact of the S355N, CONF and G376R mutations, we then visualized the corresponding residues in our THCAS 3D model including the docked CBGA (**Figure 6**). Residues S355N, CONF and G376R are located in a looped region that is adjacent to the active site and extends from Y354 to A380 (ASA-loop). This ASA-loop does not contain known catalytically active residues (**Supplemental Table S1**), but the docking demonstrates that a part of it contributes to shaping the substrate binding cavity (**Figure 6E**, blue). Within this part, residue S355 is in direct contact with the aromatic moiety and aliphatic tail of the CBGA substrate. The S355N mutation introduces an additional amine group to the sidechain, it is sterically more demanding and probably distorts the ASA-loop geometry, hampering substrate binding. This is consistent with the loss of catalytic efficiency we observed *in-vitro*. Residue CONF is located close to S355. Like S355N, the CONF mutation introduces a sterically more demanding sidechain. However, unlike S355, the CONF residue does not directly interact with the substrate (**Figure 6**). The CONF substitution therefore seems to be tolerated at this position, and its effect is not as severe as S355N. Interestingly, the CONF residue is adjacent to a loop which forms a part of the cavity that binds the substrate aliphatic tail (**Figure 6C**, **6D**). Because the aliphatic tail is shorter in CBGVA than in CBGA, its binding is probably less sterically demanding, and therefore less affected by the CONF mutation. This would explain the reduced CBGA metabolization but normal CBGVA metabolization observed for the CONF mutant (**Figure 3**). Residue G376, like CONF, does not directly interact with the substrate. The G376R mutation introduces a significantly larger side chain, which putatively distorts the loop geometry, afflicting substrate binding capability. In addition, because G376 is adjacent to K377 and K378, a third bulky residue bearing a positive charge might not be tolerated at this position. Again, this is in agreement with the observed G376R reduced activity.

**Figure 6.**
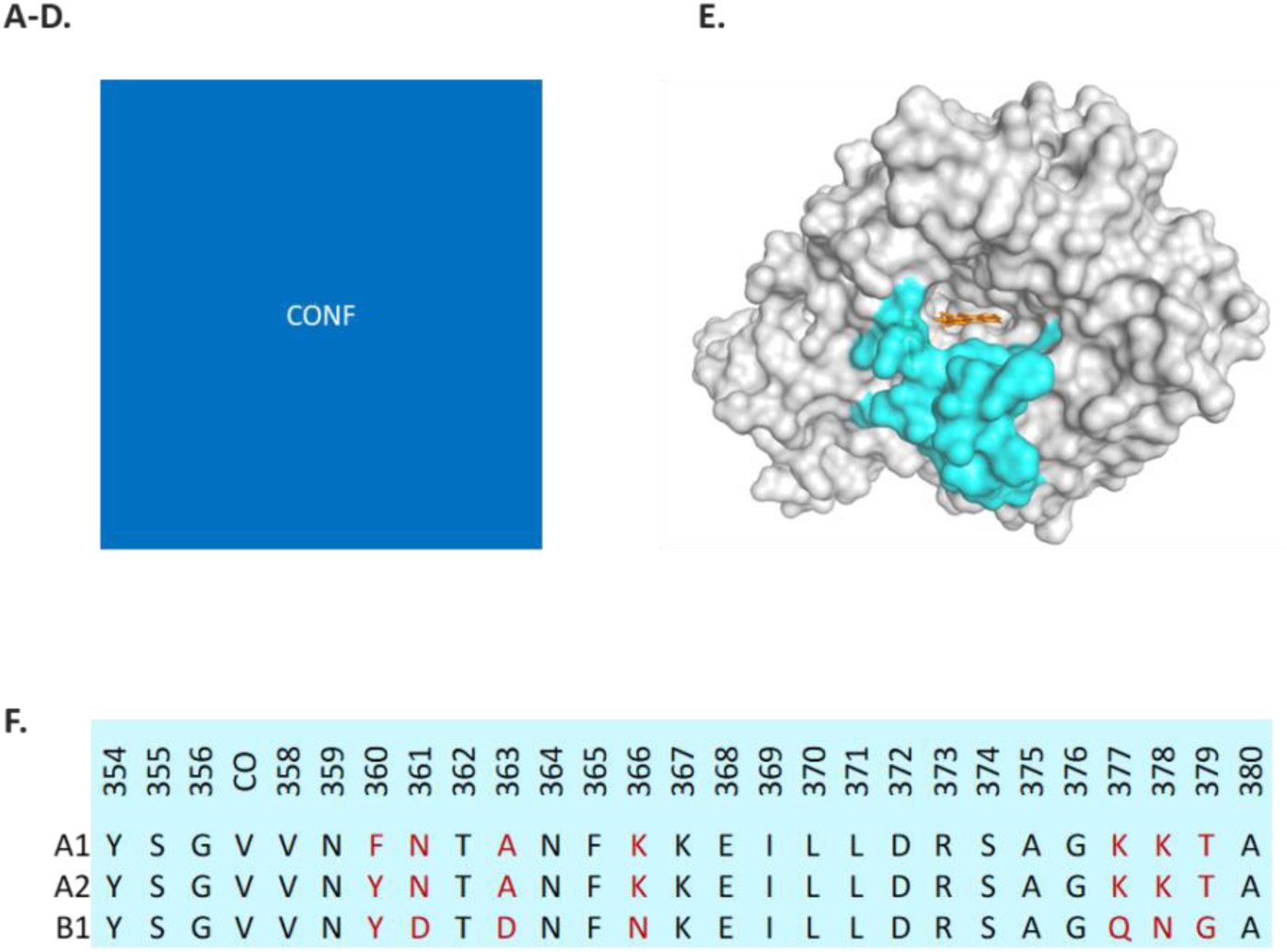
Active-site cavity of the WT THCAS and binding of CBGA in the respective cavity. **(A-D)** The ASA-loop (Y354-A380) is highlighted in pink, the respective mutations are depicted as sticks. The cavity is depicted as non-bound spheres colored by the hydrophobicity calculated for the respective coordinates with high hydrophobicity depicted in red and low in blue. The FAD cofactor is depicted as orange sticks, CBGA as green sticks. **(E)** Visualization of the ASA-loop in the THCAS enzyme. **(F)** Alignment of the ASA-loop across the THCAS (A1), CBCAS (A2) and CBDAS (B1) subclades. (**E, F**) The FAD is in orange. The enzyme is in grey, the ASA-loop in blue.

Finally, because the THCAS, CBCAS and CBDAS enzymes can all metabolize the CBGA substrate, we compared the sequence of the ASA-loop across the three enzymes (**Figure 6F**). This revealed that the ASA-loop residues are mostly conserved. In particular, the region that contributes to shaping the substrate binding cavity (**Figure 6E**), like the residues putatively responsible for the formation of Intermediate I (*i.e.*, residues Y484, Y417 and H292), are perfectly conserved. This suggests that CBGA binding and oxidation are facilitated by the same residues and mechanism in the THCAS, CBCAS and CBDAS enzymes. However, some residues of the ASA-loop are variable, primarily due to differences in the CBDAS enzyme (**Figure 6F**). Indeed, residues 361, 363, 366, 377 and 378 are lysine-rich and basic in the THCAS and CBCAS, but more acidic in the CBDAS (**Figure 6F**). Such differences in the composition of the active site putatively change the tautomerization equilibrium and the preorganization of the prenyl tail of Intermediate I and II prior to the cyclization reaction, which are different in THCAS, CBDAS and CBCAS. These variable residues may therefore contribute to the different activities observed in the three enzymes.

## Discussion

Recently, the importance of enzyme engineering has increased with the demand for enzymes adjusted to specific industrial processes. Since the number of mutation combinations that can theoretically be tested in a protein is overwhelming, a major challenge of targeted enzyme engineering consists in deciding which residues to target and how to mutate them to obtain the desired effect on the enzyme. Such targeted mutagenesis is classically guided by our structural and biochemical knowledge of the enzyme (Alejaldre et al., 2021; Victorino Da Silva Amatto et al., 2022). In this study, we successfully relied on natural variation to guide targeted mutagenesis and improve our knowledge of cannabinoid oxidocyclase mechanism. This approach had never been used before with cannabis enzymes. Our work is therefore proof of concept that studying natural variation can benefit enzyme engineering, which could be adapted to different enzymes and plant species.

### Natural variation, a treasure trove of substitutions

In many recent studies, authors investigated the natural sequence variation of cannabinoid oxidocyclase genes, aiming to link genetic variation to plant chemotype and/or identify genetic markers (Kojoma et al., 2006; McKernan et al., 2015; Onofri et al., 2015; Weiblen et al., 2015; Kitamura et al., 2016; Ali et al., 2019; Cascini et al., 2019; Doh et al., 2019; Garfinkel et al., 2021; Grassa et al., 2021; Fulvio et al., 2021; van Velzen et al., unpub.). In these studies and other cannabis genome analyses (McKernan et al., 2018; Laverty et al., 2019; McKernan et al., 2020; Gao et al., 2020), numerous cannabinoid synthase gene variants were sequenced, containing naturally occurring SNPs.

However, prior to the recent characterization of the CBCAS (Laverty et al., 2019) and establishment of a phylogenetic classification of cannabinoid oxidocyclase genes (van Velzen and Schranz, 2021), *CBCAS* genes were routinely mistaken for *THCAS* sequences. Consequently, some *THCAS* SNPs described before 2021 were in fact the WT nucleotides of their *CBCAS* homolog. This is for instance the case in Kojoma *et al*. (2006), McKernan *et al*. (2015) and Cascini *et al*. (2019). In the present study, we analyzed previously published cannabinoid oxidocyclase sequences and identified all SNPs involving aa substitution in the *THCAS*, *CBCAS* and *CBDAS* genes. The hundred mutations we list in **Supplemental Table S2** therefore represents an exhaustive and up-to-date list of aa substitutions occurring naturally in the THCAS, CBCAS and CBDAS enzymes, which can be used as a reference for future studies.

Among these mutations, some unique aa substitutions were identified in cannabis plants with unusual levels of THCA, CBDA or CBGA. The authors who identified such mutations proposed that they weaken or inactivate the associated enzymes. However, except for CONF (van Velzen et al., unpub.), authors did not perform enzyme assay to test these hypotheses (Kojoma et al., 2006; McKernan et al., 2015; Onofri et al., 2015; Garfinkel et al., 2021). From another perspective, this means that previous studies which used targeted mutagenesis (**Supplemental Table S1**) on cannabinoid oxidocyclases systematically tested artificially-designed mutations (Sirikantaramas et al., 2004; Taura et al., 2007; Shoyama et al., 2012; Zirpel et al., 2018). The only exceptions are 4 artificial mutations tested by Zirpel *et al*. (2018) which were coincidentally similar to natural substations identified somewhere else (**Table 1, Supplemental Table S1**): the artificial K377Q mutation, which has very limited impact on the THCAS (Zirpel et al., 2018), was found two years later in a *THCAS* gene variant (Gao et al. 2020). The artificial N168Q and N329Q mutations, affecting glycosylation sites (Zirpel et al., 2018), are similar to the natural N168S and N329T substitutions found in *CBDAS* and *THCAS* gene variants, respectively (Onofri et al., 2015; Cascini et al., 2019; Laverty et al., 2019; Fulvio et al., 2021). The T446I artificial mutation, which negatively impacts THCAS and CBDAS activity (Zirpel et al., 2018), is similar to the T446A substitution found in a *CBCAS* gene variant (Weiblen et al., 2015).

**Table 1.**
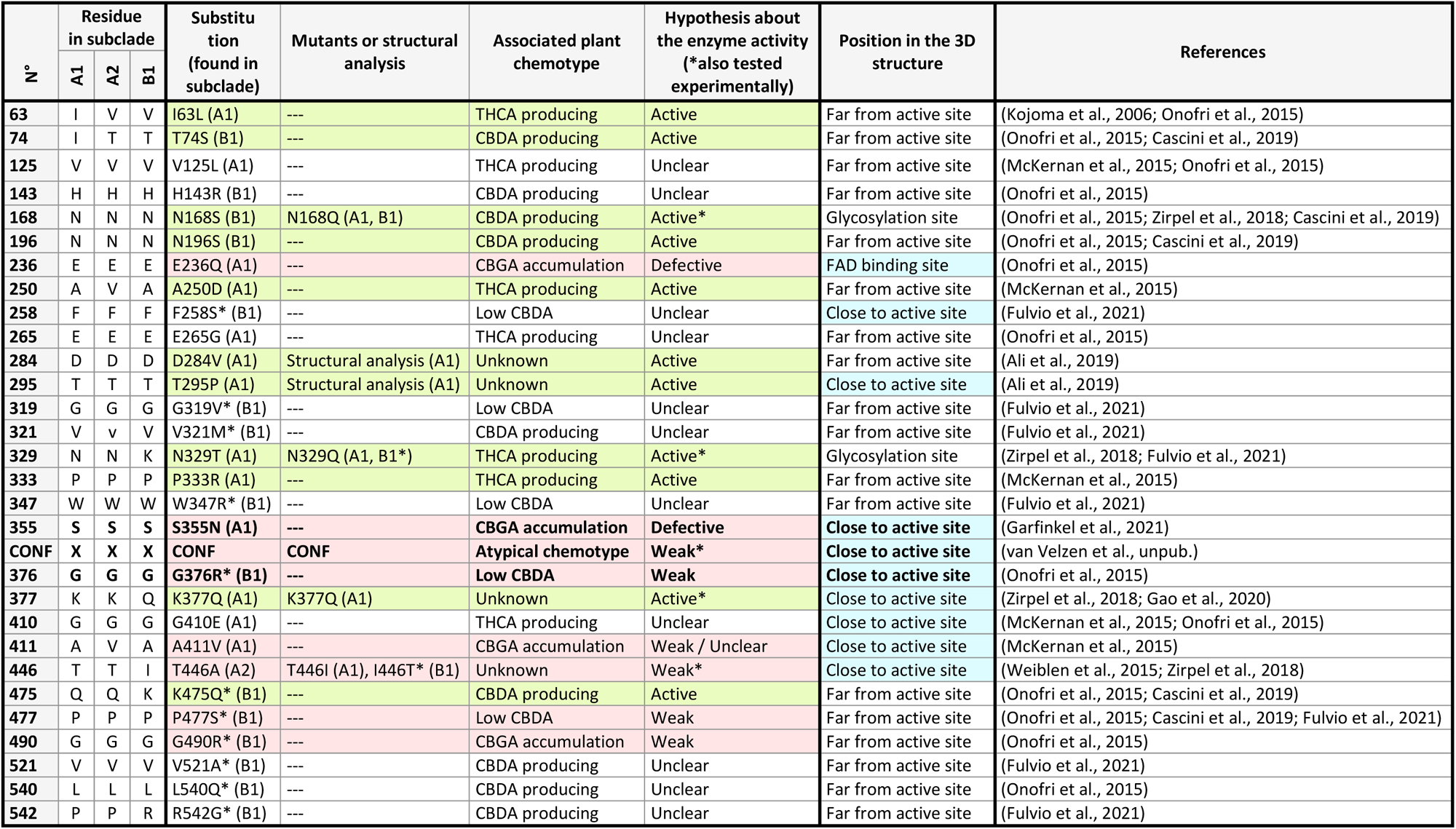
Summary of the aa substitutions associated with characterized mutants, structural analyses, and/or plant chemotype. The functional relevance of the substitutions may be experimentally measured through mutant characterization and/or hypothesized trough chemotype and structural analyses. Substitutions hypothesized or shown to have a low impact on the enzyme activity are highlighted in green. Substitutions hypothesized or shown to weaken or inactivate the enzyme are highlighted in red. The position of the substituted residues in the THCAS is detailed and the residues close to the active site or FAD binding site (i.e., the more likely to affect the enzyme activity) are highlighted in blue. The 3 substitutions of interest are highlighted in bold.

Therefore, except for CONF and the 4 artificial mutations listed above, the 95 other aa substitutions we identified in the THCAS, CBCAS and CBDAS correspond to residues that had never been investigated before. This shows how very little overlap there is between studies focusing on cannabis natural variation and the ones aiming to characterize and engineer cannabinoid oxidocyclase enzymes. To reconcile these two aspects of cannabis research, we deepened the characterization of CONF and evaluated the impact of two other naturally occurring aa substitutions found in plants with atypical chemotypes.

### S355N, CONF and G376R are responsible for plant chemotype

The S355N THCAS substitution was first described by Garfinkel *et al*. (2021), who inquired CBGA dominance inheritance in cannabis. As S355N was identified in plants with high CBGA and drastically reduced THCA levels, authors hypothesized it to inactivate the THCAS (Garfinkel et al., 2021). This substitution was later identified in several CBGA- and CBDA-dominant cannabis cultivars, suggesting it to be somehow common (McKernan et al., 2022). Our results demonstrated for the first time that the S355N THCAS mutant enzyme is unable to metabolize CBGA and only metabolize traces amounts of CBGVA. The mutation therefore severely impacts the THCAS but does not completely inactivate it. Garfinkel *et al*. found low levels of THCA in plants homozygous for the S355N mutation, which could be explained in two ways. First, the S355N mutant may metabolize trace amounts of CBGA that were below detection limit in our experiments. It is also possible that the THCA measured by Garfinkel *et al*. was synthesized as minor product of another cannabinoid synthase present in the plant. We can nonetheless conclude that the S355N substitution, by virtually inactivating the THCAS, is directly responsible for CBGA (and probably CBGVA) accumulation in cannabis.

The CONF substitution was identified in our previous study, in plants with [atypical chemotypes, CONF] (van Velzen et al., unpub.). In the present study, we demonstrated that the CONF substitution produces about two times less THCA than the WT, but as much THCVA, redirect the substrate utilization from preferential CBGA to preferential CBGVA metabolization, and alters the THCAS product specificity toward a higher production of CBC(V)A.

The G376R substitution we tested in the THCAS was first identified in the CBDAS by Onofri *et al*. (2015), who investigated the link between cannabis genotype and chemotype. In their study, authors spotted the G376R* CBDAS substitution in plants partially accumulating CBGA and producing low CBDA levels. They proposed this mutation to slightly weaken the CBDAS (Onofri et al., 2015). Our results demonstrated that the G376R THCAS mutant exhibit reduced activity, metabolizing five times less CBGA and 20 times less CBGVA than the WT. The product specificity was also impacted, as the mutant produced more CBCA and CBCVA than the WT. Our results therefore confirm that the G376R substitution significantly impairs enzymatic activity, being the cause for *in-planta* CBGA accumulation.

In summary, the S355N, CONF and G376R substitutions drastically affect cannabinoid oxidocyclase activity, in a way that explains the chemotypes of the plants in which they were identified. This means we successfully leveraged natural genetic and phenotypic variation to identify important aa residues for cannabinoid oxidocyclase activity. In the light of these results, it should therefore not be forgotten that our initial aa substitution analysis identified four other potential substitutions of interest, that may weaken or inactivate cannabinoid synthases: E236Q, A411V, P477S* and G490R* (McKernan et al., 2015; Onofri et al., 2015; Cascini et al., 2019; Fulvio et al., 2021). Residues E236 and A411 are in close proximity to the FAD binding and active site of cannabinoid oxidocyclase, making them very likely to affect enzyme activity. Residues P477* and G490* are located on the surface of the enzyme, relatively far from known regions of interest. They could therefore make original targets for future studies, with the potential to unravel another interesting part of cannabinoid oxidocyclases.

It is also interesting to note that the activity and product specificity of the WT and mutant THCAS enzymes were affected by the addition of a second substrate to the reaction mix. Indeed, compared to the incubations with CBGA or CBGVA alone, incubations performed with both substrates resulted in increased THCA/CBCA and decreased THCVA/CBCVA ratios. In a previous study, Zirpel *et al*. (2018) demonstrated that pH influences THCAS product specificity, which acid pH (4-5.5) being favorable to THCA production, and neutral pH (6.5-7.5) mostly leading to CBCA (Zirpel et al., 2018). It is therefore possible that, despite the buffer solution, reaction mixes containing CBGA and CBGVA (two cannabinoid *acids*) were more acid than the ones containing a single substrate, leading to increased THCA/CBCA ratio. This shows how sensitive cannabinoids synthases can be to small changes in their environment, putting a new perspective on the activity of these enzymes in the complex context of the cannabis plant.

### The ASA-loop; a critical region of cannabinoid oxidocyclases

In previous site-directed mutagenesis and crystallization studies, multiple residues critical for cannabinoid oxidocyclase structure and activity were identified (**Supplemental Table S1**) (Sirikantaramas et al., 2004; Taura et al., 2007; Shoyama et al., 2012; Zirpel et al., 2018). These studies contributed to a significant understanding of cannabinoid oxidocyclase structure-function relationship, helped map different regions of the enzyme (**Figure 2A**), and allowed Shoyama *et al*. (2012) to propose a catalytic mechanism for the conversion of CBGA into THCA. However, our knowledge of cannabinoid oxidocyclase enzymes is still far from being complete. It was for instance insufficient to modify the product specificity of the THCAS toward CBDA production (Zirpel et al., 2018), indicating that some critical residues or regions of cannabinoid oxidocyclases are yet to be identified. By docking the CBGA, THCA and associated reactional intermediates into the THCAS enzyme, we made it possible to visualize which THCAS residues putatively form the substrate binding pocket and interact with the CBGA substrate despite previously reported polar residues. Our results are in agreement with the impact of previously tested artificial mutations (Shoyama et al., 2012; Zirpel et al., 2018). Dockings also helped us update the THCAS catalytic mechanism proposed by Shoyama *et al*. (2012), by adding a second intermediate to the model (**Figure 5**). This helped us rationalize the impact of our mutations of interest.

The substitutions S355N, CONF and G376R affect residues that were never tested before. These residues are part of the ASA-loop, a looped region adjacent to the active site that extends from Y354 to A380. According to our bioinformatic analyses, [some residues, CONF] of the ASA-loop are highly conserved and contribute to shaping the substrate binding cavity (**Figure 6E**). The substitutions S355N, CONF and G376R, by introducing sterically more demanding sidechains in this cavity, are therefore expected to hamper substrate binding, explaining the altered activities observed in our *in-vitro* experiments. Interestingly, in a previous mutagenesis study (**Supplemental Table S1**; Zirpel et al., 2018), three other residues of the ASA-loop were tested: residues 377-379, which do not seem to shape the substrate binding cavity. As residues 377-379 are different in the THCAS/CBCAS (K377, K378, T379) and CBDAS (Q377*, N378*, G379*) enzymes (**Figure 6F**), Zirpel *et al*. (2018) generated a triple THCAS-mutant enzyme possessing the three CBDAS equivalent (THCAS-K377Q, K378N, T379G). This triple mutant retained 88% of the activity of the WT, demonstrating limited impact of the mutations (Zirpel et al., 2018). Taken together, we can therefore separate the ASA-loop into two regions: on the one hand, [some residues, CONF] form a region that is highly conserved across cannabinoid oxidocyclases, and putatively involved in substrate binding. Even a rather small change in this region can have a significant impact on substrate metabolization. This emphasizes the sensitivity of this region towards mutations and its critical role in enzyme activity, suggesting that mutating other residues in this region could similarly alter the activity of the THCAS, CBCAS and CBDAS enzymes. On the other hand, the remaining residues of the ASA-loop are more variable across cannabinoid oxidocyclases. They are unlikely to be involved in substrate binding and are not as critical for the enzyme activity, but they may impact product specificity.

To conclude, by functionally characterizing the impact of S355N, CONF and G376R, 3 naturally occurring SNPs, we highlighted a new critical part of cannabinoid oxidocyclase enzymes. Our study demonstrates how natural variation can help decipher the mechanism and properties of an enzyme which, in turn, as an example of reciprocal illumination, may help guide breeding and the development of cultivars with specific cannabinoid profiles.

## Material and methods

### Material

Standard solutions of cannabinoids 1.0 mg mL^-1^ in acetonitrile were purchased from Sigma Aldrich.

### Aa substitutions analysis

A cannabinoid oxidocyclase nucleotide sequence dataset was constituted by adding sequences from McGarvey *et al*. (2020), Fulvio *et al*. (2021) and Garfinkel *et al*. (2021) to the recent van Velzen and Schranz (2021) compilation. Most pseudogenes and ambiguous sequences (*e.g.*, potential chimeras) were excluded, apart from a few A3, A4 and B2 pseudogenes, which were kept to retain sequences representative of every subclade. Sequences were aligned with MAFFT v7.490, using default parameters (Katoh et al., 2019), and manually curated. A maximum likelihood gene tree was built with IQ-TREE, using the automatic substitution model selection and a bootstrap alignments number of 5000 (Trifinopoulos et al., 2016). The tree was visualized with FIGTREE v.1.4.4. Aligned nucleotide sequences from the A1, A2 and B1 subclades were retrieved to form three sub-datasets, which were translated into aa and manually analyzed in Bioedit (Hall, 1999) to identify SNPs involving aa substitutions in the THCAS, CBCAS and CBDAS enzymes, respectively. Residues impacted by the substitutions were visualized on the THCAS crystal structure (PDB ID: 3vte.1.A) (Shoyama et al., 2012) using PyMOL (v.2.5.2, Schrödinger, LLC). All aa substitutions and associated data were reported in **Supplemental Table S1**.

### Synthesis and cloning of the *THCAS* genes

Coding sequences of the WT *THCAS* (GenBank accession number AB057805) and associated mutants S355N, CONF and G376R were synthesized and subcloned into the pPICZαA expression vector (Invitrogen™, Thermo Fisher Scientific) by GenScript® (Leiden, the Netherlands. Specifically, sequences were synthesized without their native Cannabis signal peptide, a 6xHis tag extension was added at the 3’ end, internal *Eco*RI and *Xba*I restriction sites were removed using genetic code redundancy, and the sequences were framed by *Eco*RI and *Xba*I restriction sites at the 5’ and 3’ ends, respectively. Synthetic sequences were then subcloned into pPICZαA using *Eco*RI and *Xba*I restriction enzymes, in frame with the plasmid N-terminal α-factor.

### Expression and purification of the THCAS enzyme

Recombinant pPICZαA plasmids containing the WT and mutant *THCAS* were used to transform the *Komagataella phaffii* strain X-33 (Invitrogen™, Thermo Fisher Scientific), as specified by the supplier. THCAS heterologous expression was achieved by culturing the recombinant *K. phaffi* according to the supplier’s recommendations, with modifications based on Taura *et al*. (2007) and Karbalaei *et al*. (2020). Briefly, an isolated colony of recombinant *K. phaffi* was picked into 100 mL of classic BMGY culture medium (10 g L^-1^ yeast extract, 20 g L^-1^ peptone, 1.34% yeast nitrogen base, 4 × 10^-5^% biotin, 1% glycerol, 100 mM potassium phosphate buffer pH 6.0) and cultured at 30 °C, 230 rpm for 16-18 h. The yeast suspension was centrifuged at 4,000 x g for 5 min and the pellet was resuspended at a cell density corresponding to OD_600_ 1.0 in 50 mL modified BMMY induction medium (10 g.L^-1^ yeast extract, 20 g.L^-1^ peptone, 5 g.L^-1^ bacto casamino acids, 10 g.L^-1^ sorbitol, 1.34% yeast nitrogen base, 4 × 10^-5^ % biotin, 0.001% riboflavin, 1% methanol, 100 mM sodium citrate buffer pH 5.5). The yeast suspension was incubated at 30°C and 230 rpm for 48 h, with the addition of 0.5% methanol after the first 24 h to maintain the induction. It is important to note that *K. phaffi* is normally cultured in baffled flasks, but optimal THCAS production was achieved with non-baffled flasks.

After 48h of induction, the 50 mL culture supernatant containing the soluble THCAS was collected by centrifugation at 4000 x g, 4 °C for 10 min. The enzyme solution was concentrated and buffer-exchanged into ≈ 2 mL of binding buffer (20 mM sodium phosphate, 500 mM NaCl, 20 mM imidazole, pH 7.4) with a 10 kDA cut-off Amicon® Ultra-15 Centrifugal Filter Unit (Millipore). The concentrated solution was applied onto a His SpinTrap™ column (Cytiva) for purification of the His-tagged THCAS. Sample application and washing were performed in native conditions, as described by the supplier, but elution was performed in modified elution buffer (20 mM sodium citrate, 500 mM NaCl, 500 mM imidazole, pH 5.5). Finally, the purified THCAS solution was buffer-exchanged into ≈ 40 µL of 100 mM sodium citrate (pH 5.0) with a 10 kDA cut-off Amicon® Ultra-0.5 Centrifugal Filter Unit (Millipore). Enzyme was kept on ice and used fresh for enzyme assay. The presence of the His-tagged THCAS in the purified solution was verified by Western-blot; the enzyme concentration was measured with a Bradford test.

### THCAS *in-vitro* assays

To determine if the enzymes were active, qualitative assays were performed by incubating 20-40 µg.mL^-1^ of freshly produced enzymes in 50 µL of 100 mM sodium citrate buffer (pH 5.0) containing 100 µM of either CBGA or CBGVA. After 30 min at 45°C, 700 rpm, reactions were stopped by the addition of 1 volume acetonitrile 100%.

Because of cannabinoids have low solubility in aqueous buffers (Zirpel et al., 2015), assays had to be done with substrate concentrations <100-150 µM. In preliminary experiments, incubations with CBGA concentrations ranging from 5 to 100 µM only yielded a linear part of the Michaelis–Menten curve (data not shown), which is consistent with a Km > 100 µM (Taura et al. 1995). In these conditions, the kinetic parameters of the enzymes could not be accurately determined. Therefore, to compare the performances of the WT and mutants THCAS, standardized assays were performed by incubating 15 µg.mL^-1^ of fresh enzymes in 50 µL of 100 mM sodium citrate buffer (pH 5.0) containing 50 µM of CBGA and/or CBGVA. After 10 min at 25°C, 700 rpm, reactions were stopped by the addition of 1 volume acetonitrile 100%. Reactions were performed in triplicates.

### Identification and quantification of cannabinoids

Reaction mixtures were filtered at 0.2 μm and analyzed using an ultra-high performance liquid chromatography (UHPLC) UltiMate™ 3000 (Thermo Scientific), equipped with a LiChrospher® 100 RP-18 (125 mm x 4 mm, 5 µm) C18 reversed-phase column (Merck). The elution solvents consisted of ultrapure water with 0.1% formic acid (A) and acetonitrile (B). The mobile gradient phase was as follows (A/B; v/v): 60 : 40 between 0 and 1 min, 30 : 70 at 4 min, 10 : 90 at 19 min, 0 : 100 at 20 min, and 60 : 40 between 25 and 30 min. Samples were injected with a volume of 20 μL. During the runtime, the mobile phase flow rate was kept at 0.8 mL.min^-1^ and the column at 40°C. Compounds were detected based on UV scans at 270 nm. Data were recorded and analyzed on Chromeleon ™ (v.7.2.10, Thermo Scientific). Reaction products were identified by comparison of their retention time with those of standard molecules.

### *In-silico* docking experiments

Molecular docking experiments were performed to identify a putative productive binding mode for the substrate CBGA and a binding mode of the product THCA in THCAS. Molecular docking was performed using YASARA embedded in the CavitomiX™ platform (Innophore, Graz, Austria) (Gruber et al., 2020) using the catalophore point cloud technology for cavity guided docking (Krieger et al., 2002; Hetmann et al., 2023). As template the crystal structure of the WT THCAS from *Cannabis sativa*, crystalized by Shoyama *et al*. (2012) (PDB ID: 3vte) was used. The structures of CBGA and THCA as well as the missing loops of the crystal structure of THCAS were built utilizing Yasara (Vers. 22.9.24.W.64). First, the physico-chemical properties inside the cavity were analyzed and represented as 3D point clouds. Following settings were used: Min cav. Vol: 300 Å³; max cav. Vol: 1500 Å³; ligsite cutoff 5; probe radius 1 Å; Grid Spacing 0.375; Softshell 0.4 Å. The docking experiment was performed using following settings: Docking box extension around cavity: 2Å; Docking runs: 5; Clustering RMSD: 1 Å. The computed results were visualized, analyzed and the figures were prepared using PyMOL™ (Vers. 2.5.4, Schrödinger, LLC).

## Acknowledgments

The authors would like to thank Ludivine Hocq, Marwa Roumani and Alain Hehn for advice on molecular biology and enzymology.

## Author contributions

CV, BD, and RvV designed the experiments. CV and NPM conducted the *in-vitro* experiments, with support of ACvdP, FV, KC and IFK. CV, CB, and BD performed the *in-silico* analyses. CV, CB, NP and BD analysed the data. CV, MES, BD and RvV wrote the article. MES, BD, and RvV directed the project.

No conflict of interest declared.

## References

Alejaldre, L., Pelletier, J. N., and Quaglia, D. (2021). Methods for enzyme library creation: Which one will you choose?: A guide for novices and experts to introduce genetic diversity. BioEssays 43:2100052.

Ali, S., Mufti, M., Khan, M., and Aziz, I. (2019). The identification of SNPs in THCA synthase gene from Pakistani Cannabis. Asia Pac J Mol Biol Biotechnol 27:1–9.

Boggs, D. L., Nguyen, J. D., Morgenson, D., Taffe, M. A., and Ranganathan, M. (2018). Clinical and Preclinical Evidence for Functional Interactions of Cannabidiol and Δ9-Tetrahydrocannabinol. Neuropsychopharmacology 43:142–154.

Burstein, S. (2015). Cannabidiol (CBD) and its analogs: a review of their effects on inflammation. Bioorg. Med. Chem. 23:1377–1385.

Casajuana Köguel, C., López-Pelayo, H., Balcells-Olivero, M. M., Colom, J., and Gual, A. (2018). Psychoactive constituents of cannabis and their clinical implications: a systematic review. Adicciones 30:140–151.

Cascini, F., Farcomeni, A., Migliorini, D., Baldassarri, L., Boschi, I., Martello, S., Amaducci, S., Lucini, L., and Bernardi, J. (2019). Highly Predictive Genetic Markers Distinguish Drug-Type from Fiber-Type Cannabis sativa L. Plants 8:496.

Colizzi, M., Ruggeri, M., and Bhattacharyya, S. (2020). Unraveling the Intoxicating and Therapeutic Effects of Cannabis Ingredients on Psychosis and Cognition. Front. Psychol. 11:833.

Costa, B. (2007). On the Pharmacological Properties of Δ9-Tetrahydrocannabinol (THC). Chem. Biodivers. 4:1664–1677.

Daniel, B., Konrad, B., Toplak, M., Lahham, M., Messenlehner, J., Winkler, A., and Macheroux, P. (2017). The family of berberine bridge enzyme-like enzymes: A treasure-trove of oxidative reactions. Arch. Biochem. Biophys. 632:88–103.

De Meijer, E. P. M., and Hammond, K. M. (2016). The inheritance of chemical phenotype in Cannabis sativa L. (V): regulation of the propyl-/pentyl cannabinoid ratio, completion of a genetic model. Euphytica 210:291–307.

Doh, E. J., Lee, G., Yun, Y.-J., Kang, L.-W., Kim, E. S., Lee, M. Y., and Oh, S.-E. (2019). DNA Markers to Discriminate Cannabis sativa L. ‘Cheungsam’ with Low Tetrahydrocannabinol (THC) Content from Other South Korea Cultivars Based on the Nucleotide Sequences of Tetrahydrocannabinolic Acid Synthase and Putative 3-Ketoacyl-CoA Synthase Genes. Evid Based Complement Altern. Med 2019:1–10.

ElSohly, M. A., Radwan, M. M., Gul, W., Chandra, S., and Galal, A. (2017). Phytochemistry of Cannabis sativa L. In Phytocannabinoids (ed. Kinghorn, A. D.), Falk, H.), Gibbons, S.), and Kobayashi, J.), pp. 1–36. Cham: Springer International Publishing.

Fellermeier, M., and Zenk, M. H. (1998). Prenylation of olivetolate by a hemp transferase yields cannabigerolic acid, the precursor of tetrahydrocannabinol. FEBS Lett. 427:283–285.

Fraaije, M. W., and Mattevi, A. (2000). Flavoenzymes: diverse catalysts with recurrent features. Trends Biochem. Sci. 25:126–132.

Fulvio, F., Paris, R., Montanari, M., Citti, C., Cilento, V., Bassolino, L., Moschella, A., Alberti, I., Pecchioni, N., Cannazza, G., et al. (2021). Analysis of Sequence Variability and Transcriptional Profile of Cannabinoid synthase Genes in Cannabis sativa L. Chemotypes with a Focus on Cannabichromenic acid synthase. Plants 10:1857.

Gagne, S. J., Stout, J. M., Liu, E., Boubakir, Z., Clark, S. M., and Page, J. E. (2012). Identification of olivetolic acid cyclase from Cannabis sativa reveals a unique catalytic route to plant polyketides. Proc. Natl. Acad. Sci. 109:12811–12816.

Gao, S., Wang, B., Xie, S., Xu, X., Zhang, J., Pei, L., Yu, Y., Yang, W., and Zhang, Y. (2020). A high-quality reference genome of wild Cannabis sativa. Hortic. Res. 7:73.

Gaoni, Y., and Mechoulam, R. (1971). Isolation and structure of .DELTA.+-tetrahydrocannabinol and other neutral cannabinoids from hashish. J. Am. Chem. Soc. 93:217–224.

Garfinkel, A. R., Otten, M., and Crawford, S. (2021). SNP in Potentially Defunct Tetrahydrocannabinolic Acid Synthase Is a Marker for Cannabigerolic Acid Dominance in Cannabis sativa L. Genes 12:228.

Grassa, C. J., Weiblen, G. D., Wenger, J. P., Dabney, C., Poplawski, S. G., Motley, S. T., Michael, T. P., and Schwartz, C. J. (2021). A new *Cannabis* genome assembly associates elevated cannabidiol (CBD) with hemp introgressed into marijuana. New Phytol. 230:1665–1679.

Gruber, K., Steinkellner, G., and Gruber, C. (2020). Determining novel enzymatic functionalities using three-dimensional point clouds representing physico chemical properties of protein cavities Advance Access published 2020.

Hall, T. A. (1999). BioEdit: a user-friendly biological sequence alignment editor and analysis program for Windows 95/98/NT. Nucleic Acids Symp. Ser. 41:95–98.

Hetmann, M., Langner, C., Durmaz, V., Cespugli, M., Köchl, K., Krassnigg, A., Blaschitz, K., Groiss, S., Loibner, M., Ruau, D., et al. (2023). Identification and validation of fusidic acid and flufenamic acid as inhibitors of SARS-CoV-2 replication using DrugSolver CavitomiX. Sci. Rep. 13:11783.

Karbalaei, M., Rezaee, S. A., and Farsiani, H. (2020). *Pichia pastoris*: A highly successful expression system for optimal synthesis of heterologous proteins. J. Cell. Physiol. 235:5867–5881.

Katoh, K., Rozewicki, J., and Yamada, K. D. (2019). MAFFT online service: multiple sequence alignment, interactive sequence choice and visualization. Brief. Bioinform. 20:1160–1166.

Kitamura, M., Aragane, M., Nakamura, K., Watanabe, K., and Sasaki, Y. (2016). Development of Loop-Mediated Isothermal Amplification (LAMP) Assay for Rapid Detection of *Cannabis sativa*. Biol. Pharm. Bull. 39:1144–1149.

Kojoma, M., Seki, H., Yoshida, S., and Muranaka, T. (2006). DNA polymorphisms in the tetrahydrocannabinolic acid (THCA) synthase gene in “drug-type” and “fiber-type” Cannabis sativa L. Forensic Sci. Int. 159:132–140.

Krieger, E., Koraimann, G., and Vriend, G. (2002). Increasing the precision of comparative models with YASARA NOVA-a self-parameterizing force field. Proteins Struct. Funct. Bioinforma. 47:393–402.

Laverty, K. U., Stout, J. M., Sullivan, M. J., Shah, H., Gill, N., Holbrook, L., Deikus, G., Sebra, R., Hughes, T. R., Page, J. E., et al. (2019). A physical and genetic map of *Cannabis sativa* identifies extensive rearrangements at the *THC/CBD acid synthase* loci. Genome Res. 29:146–156.

Livingston, S. J., Quilichini, T. D., Booth, J. K., Wong, D. C. J., Rensing, K. H., Laflamme-Yonkman, J., Castellarin, S. D., Bohlmann, J., Page, J. E., and Samuels, A. L. (2020). Cannabis glandular trichomes alter morphology and metabolite content during flower maturation. Plant J. 101:37–56.

Luo, X., Reiter, M. A., d’Espaux, L., Wong, J., Denby, C. M., Lechner, A., Zhang, Y., Grzybowski, A. T., Harth, S., Lin, W., et al. (2019). Complete biosynthesis of cannabinoids and their unnatural analogues in yeast. Nature 567:123–126.

Lupica, C. R., Riegel, A. C., and Hoffman, A. F. (2004). Marijuana and cannabinoid regulation of brain reward circuits: Marijuana and central reward circuits. Br. J. Pharmacol. 143:227–234.

McKernan, K. J., Helbert, Y., Tadigotla, V., McLaughlin, S., Spangler, J., Zhang, L., and Smith, D. (2015). Single molecule sequencing of THCA synthase reveals copy number variation in modern drug-type Cannabis sativa L. BioRxiv 028654.

McKernan, K., Helbert, Y., Kane, L. T., Ebling, H., Zhang, L., Liu, B., Eaton, Z., Sun, L., Dimalanta, E. T., Kingan, S., et al. (2018). Cryptocurrencies and Zero Mode Wave guides: An unclouded path to a more contiguous Cannabis sativa L. genome assembly. Open Science Framework.

McKernan, K. J., Helbert, Y., Kane, L. T., Ebling, H., Zhang, L., Eaton, Z., McLaughlin, S., Kingan, S., Baybayan, P., Jordan, M., et al. (2020). Sequence and annotation of 42 cannabis genomes reveals extensive copy number variation in cannabinoid synthesis and pathogen resistance genes. BioRxiv 894428.

McKernan, K., Zhang, L., Helbert, Y., Kane, L. T., and McLaughlin, S. (2022). Missense mutations in THCAS are associated with Cannabigerolic Acid expression in Cannabis sativa L. *Zenodo* Advance Access published January 25, 2022, doi:10.5281/ZENODO.63CONF30.

Mechoulam, R. (2005). Plant cannabinoids: a neglected pharmacological treasure trove: Commentary. Br. J. Pharmacol. 146:913–915.

Morimoto, S., Komatsu, K., Taura, F., and Shoyama, Y. (1998). Purification and characterization of cannabichromenic acid synthase from Cannabis sativa. Phytochemistry 49:1525–1529.

Onofri, C., de Meijer, E. P. M., and Mandolino, G. (2015). Sequence heterogeneity of cannabidiolic- and tetrahydrocannabinolic acid-synthase in Cannabis sativa L. and its relationship with chemical phenotype. Phytochemistry 116:57–68.

Shoyama, Y., Tamada, T., Kurihara, K., Takeuchi, A., Taura, F., Arai, S., Blaber, M., Shoyama, Y., Morimoto, S., and Kuroki, R. (2012). Structure and Function of Δ1-Tetrahydrocannabinolic Acid (THCA) Synthase, the Enzyme Controlling the Psychoactivity of Cannabis sativa. J. Mol. Biol. 423:96–105.

Sirikantaramas, S., Morimoto, S., Shoyama, Y., Ishikawa, Y., Wada, Y., Shoyama, Y., and Taura, F. (2004). The Gene Controlling Marijuana Psychoactivity. J. Biol. Chem. 279:39767–39774.

Sirikantaramas, S., Taura, F., Tanaka, Y., Ishikawa, Y., Morimoto, S., and Shoyama, Y. (2005). Tetrahydrocannabinolic Acid Synthase, the Enzyme Controlling Marijuana Psychoactivity, is Secreted into the Storage Cavity of the Glandular Trichomes. Plant Cell Physiol. 46:1578–1582.

Stout, J. M., Boubakir, Z., Ambrose, S. J., Purves, R. W., and Page, J. E. (2012). The hexanoyl-CoA precursor for cannabinoid biosynthesis is formed by an acyl-activating enzyme in Cannabis sativa trichomes: A cytoplasmic acyl-activating enzyme involved in cannabinoid biosynthesis. Plant J. 71:353– 365.

Suraev, A. S., Marshall, N. S., Vandrey, R., McCartney, D., Benson, M. J., McGregor, I. S., Grunstein, R. R., and Hoyos, C. M. (2020). Cannabinoid therapies in the management of sleep disorders: A systematic review of preclinical and clinical studies. Sleep Med. Rev. 53:101339.

Taura, F., Morimoto, S., Shoyama, Y., and Mechoulam, R. (1995). First direct evidence for the mechanism of D1-tetrahydrocannabinolic acid biosynthesis. J. Am. Chem. Soc. 117:9766–9767.

Taura, F., Morimoto, S., and Shoyama, Y. (1996). Purification and Characterization of Cannabidiolic-acid Synthase from Cannabis sativa L. J. Biol. Chem. 271:17411–17416.

Taura, F., Sirikantaramas, S., Shoyama, Y., Yoshikai, K., Shoyama, Y., and Morimoto, S. (2007). Cannabidiolic-acid synthase, the chemotype-determining enzyme in the fiber-type *Cannabis sativa*. FEBS Lett. 581:2929–2934.

Taura, F., Tanaka, S., Taguchi, C., Fukamizu, T., Tanaka, H., Shoyama, Y., and Morimoto, S. (2009). Characterization of olivetol synthase, a polyketide synthase putatively involved in cannabinoid biosynthetic pathway. FEBS Lett. 583:2061–2066.

Trifinopoulos, J., Nguyen, L.-T., von Haeseler, A., and Minh, B. Q. (2016). W-IQ-TREE: a fast online phylogenetic tool for maximum likelihood analysis. Nucleic Acids Res. 44:W232–W235.

van de Donk, T., Niesters, M., Kowal, M. A., Olofsen, E., Dahan, A., and van Velzen, M. (2019). An experimental randomized study on the analgesic effects of pharmaceutical-grade cannabis in chronic pain patients with fibromyalgia. Pain 160:860–869.

van Velzen, R., and Schranz, M. E. (2021). Origin and Evolution of the Cannabinoid Oxidocyclase Gene Family. Genome Biol. Evol. 13:evab130.

Victorino Da Silva Amatto, I., Gonsales Da Rosa-Garzon, N., Antônio De Oliveira Simões, F., Santiago, F., Pereira Da Silva Leite, N., Raspante Martins, J., and Cabral, H. (2022). Enzyme engineering and its industrial applications. Biotechnol. Appl. Biochem. 69:389–409.

Wang, M., Wang, Y.-H., Avula, B., Radwan, M. M., Wanas, A. S., van Antwerp, J., Parcher, J. F., ElSohly, M. A., and Khan, I. A. (2016). Decarboxylation Study of Acidic Cannabinoids: A Novel Approach Using Ultra-High-Performance Supercritical Fluid Chromatography/Photodiode Array-Mass Spectrometry. Cannabis Cannabinoid Res. 1:262–271.

Weiblen, G. D., Wenger, J. P., Craft, K. J., ElSohly, M. A., Mehmedic, Z., Treiber, E. L., and Marks, M. D. (2015). Gene duplication and divergence affecting drug content in *Cannabis sativa*. New Phytol. 208:1241–1250.

Winkler, A., Motz, K., Riedl, S., Puhl, M., Macheroux, P., and Gruber, K. (2009). Structural and Mechanistic Studies Reveal the Functional Role of Bicovalent Flavinylation in Berberine Bridge Enzyme. J. Biol. Chem. 284:19993–20001.

Zirpel, B., Stehle, F., and Kayser, O. (2015). Production of Δ9-tetrahydrocannabinolic acid from cannabigerolic acid by whole cells of Pichia (Komagataella) pastoris expressing Δ9-tetrahydrocannabinolic acid synthase from Cannabis sativa l. Biotechnol. Lett. 37:1869–1875.

Zirpel, B., Kayser, O., and Stehle, F. (2018). Elucidation of structure-function relationship of THCA and CBDA synthase from Cannabis sativa L. J. Biotechnol. 284:17–26.

